# Blood-spinal cord barrier leakage is independent of motor neuron pathology in ALS

**DOI:** 10.1101/704270

**Authors:** Sarah Waters, Birger V. Dieriks, Molly E. V. Swanson, Yibin Zhang, Natasha L. Grimsey, Helen C. Murray, Clinton Turner, Henry J. Waldvogel, Richard L.M. Faull, Jiyan An, Robert Bowser, Maurice A. Curtis, Mike Dragunow, Emma L. Scotter

**Author notes:** Corresponding authors Emma Scotter. Mike Dragunow.

## Abstract

**Background:** Amyotrophic lateral sclerosis (ALS) is a fatal neurodegenerative disease involving progressive degeneration of upper and lower motor neurons. Both lower motor neuron loss and the deposition of phosphorylated TDP-43 inclusions display regional patterning along the spinal cord. The blood-spinal cord barrier (BSCB) ordinarily restricts entry into the spinal cord parenchyma of blood components that are neurotoxic, but in ALS there is evidence for barrier breakdown. Here we sought to examine whether BSCB breakdown, motor neuron loss, and TDP-43 proteinopathy display the same regional patterning across and along the spinal cord.

**Methods:** We measured cerebrospinal fluid (CSF) hemoglobin in living ALS patients (n=87 controls, n=236 ALS) as a potential biomarker of BSCB and blood-brain barrier leakage. We then immunostained cervical, thoracic, and lumbar post mortem spinal cord tissue (n=5 controls, n=13 ALS) and employed semi-automated imaging and analysis to quantify and map lower motor neuron loss and phosphorylated TDP-43 inclusion load against hemoglobin leakage.

**Results:** Motor neuron loss and TDP-43 proteinopathy were seen at all three levels of the ALS spinal cord, with most abundant TDP-43 deposition in the ventral grey (lamina IX) of the cervical and lumbar cord. In contrast, hemoglobin leakage was observed along the ALS spinal cord axis but was most severe in the dorsal grey and white matter in the thoracic spinal cord.

**Conclusions:** Our data show that leakage of the blood-spinal cord barrier occurs during life but at end-stage its distribution is independent from the major motor neuron pathology and is unlikely to be a major contributor to pathogenesis in ALS.

## Background

Amyotrophic lateral sclerosis (ALS) is characterised by the progressive and eventually fatal degeneration of upper and lower motor neurons [1–3]. Motor symptom onset is focal, beginning most commonly in the limbs, less commonly in the muscles of speech and swallowing, and rarely in the respiratory muscles [1–3]. Recently it was shown that the motor neurons that innervate the area of first symptom onset show the greatest cell loss and pathologic deposition of TDP-43 [4, 5]. A wealth of evidence shows that cell-autonomous TDP-43 dysfunction initiates motor neuron degeneration [6–8], and indeed phosphorylated TDP-43 protein inclusions are seen in almost all ALS cases [9, 10]. However, degeneration can also be modified by non-cell autonomous factors including glial activity and the integrity of the blood-brain and blood-spinal cord barriers [8, 11].

The blood-brain barrier (BBB), blood-spinal cord barrier (BSCB), and blood-cerebrospinal fluid (blood-CSF) barrier are specialised interfaces that regulate the influx of nutrients and ions into the central nervous system, and the removal of waste and other solutes [12, 13]. In addition, these barriers separate the brain, spinal cord parenchyma, and CSF respectively from potentially neurotoxic blood-borne components in the circulation, such as peripheral leukocytes, erythrocytes and plasma proteins [14–16]. The blood-CSF barrier is formed by apical tight junctions between the epithelial cells of the choroid plexus, which prevent the passage of solutes from the blood to the CSF [13]. The choroid plexus is also the primary producer of the CSF [17], however around one-third of the CSF is derived from the CNS parenchymal interstitial fluid [18]. In contrast, the BBB and BSCB comprise endothelial cells knitted together by tight junctions, with a basement membrane of structural matrix proteins, pericytes embedded within this matrix, and astrocytes whose endfeet encircle the basement membrane [19].

In ALS, the BBB and BSCB are compromised. Their disruption is evidenced by extravasation of blood components such as immunoglobulin (IgG), fibrin, thrombin and the erythrocyte components hemoglobin and hemosiderin into the brain and spinal cord parenchyma [20, 21]. Blood protein concentrations are also altered in ALS CSF, implicating either blood-CSF barrier disruption, and/or diffusion of BBB and BSCB leakage factors from the brain and spinal cord interstitial fluid into the CSF [22]. Many of these blood components are neurotoxins [23–27] which, when allowed into contact with motor neurons, are likely to enhance motor neuron vulnerability to autonomous TDP-43-mediated damage.

Autonomous motor neuron damage and death in most cases of ALS is driven by misfolding, mislocalisation and loss of RNA processing function of TDP-43 [6, 28]. Indeed, the regional distribution of neuronal TDP-43 protein inclusions across the brain and spinal cord is positively correlated with the degree of cell death, implicating TDP-43 misfolding and deposition as a key mechanistic event [4, 5, 29]. In the spinal cord, both TDP-43 proteinopathy and neuronal loss are greatest at the cervical and lumbar enlargements where their severity correlates with the site of symptom onset [5]. The patterning along the spinal cord axis provides a well-characterised setting against which to explore the integrity of the BSCB.

Here we investigate the spatial relationship between BSCB compromise, TDP-43 proteinopathy and motor neuron loss along the spinal cord, in order to determine whether BSCB leakage may be a contributor to human ALS pathogenesis and thus a target for therapy.

## Methods

### CSF samples and patient spinal cord tissue

Cerebrospinal fluid samples provided by the NEALS Biofluid Repository, California, USA, were obtained via lumbar puncture, upon informed patient consent (Table 1). ALS diagnosis was performed by licensed neurologists specialized in motor neuron disease, using revised El Escorial criteria. None of the samples exhibited visible blood contamination. Samples were spun at 3000 rpm at 4 °C for 10 min to remove cells and debris, then aliquoted and stored in low protein-binding polypropylene tubes at −80 °C within 2 h of collection.

**Table 1.** CSF sample donor demographics.

|  | Healthy Controls<br>(n = 87) | ALS<br>(n = 236) |
| --- | --- | --- |
| Age at donation<br>(mean ± SD) | 48.4 ± 16.3 | 53.7 ± 14.2 |
| Sex (M / F) | 42 / 45 | 166 / 70 |
| Disease Type<br>(Sporadic / Familial) | N/A | 205 / 31 |

Post-mortem human spinal cord tissue from were obtained from the New Zealand Neurological Foundation Human Brain Bank at the Centre for Brain Research, Auckland, New Zealand (Table 2). Consent was obtained from donors and their families prior to death and ethical approval for this study was obtained from The University of Auckland Human Participants Ethics Committee. Clinical diagnosis of ALS was performed by consultant neurologists at Auckland City or Middlemore Hospitals, Auckland, New Zealand. Neuropathological diagnosis was performed by consultant neuropathologists at Auckland City Hospital. Control spinal cords were obtained from cadavers fixed by injection of anatomical embalming fluid (Dodge Anatomical mix) via the carotid artery; ALS cords were immersion fixed in 15% formalin. Three spinal cord regions were analysed: cervical (C8), mid-thoracic (T7-T9), and lumbar (L4/L5). To select the appropriate segmental block for each case we first measured the transverse diameters of all spinal cord segments with Vernier callipers (Fig S1A) and compared them with population estimates of segmental diameter (Fig S1B) [30]. This demonstrated the disparity between cords sectioned and labelled according to vertebral level compared to neuronal level. Spinal cord segments were then relabelled for neuronal level using a conversion scheme [30].

**Table 2.**
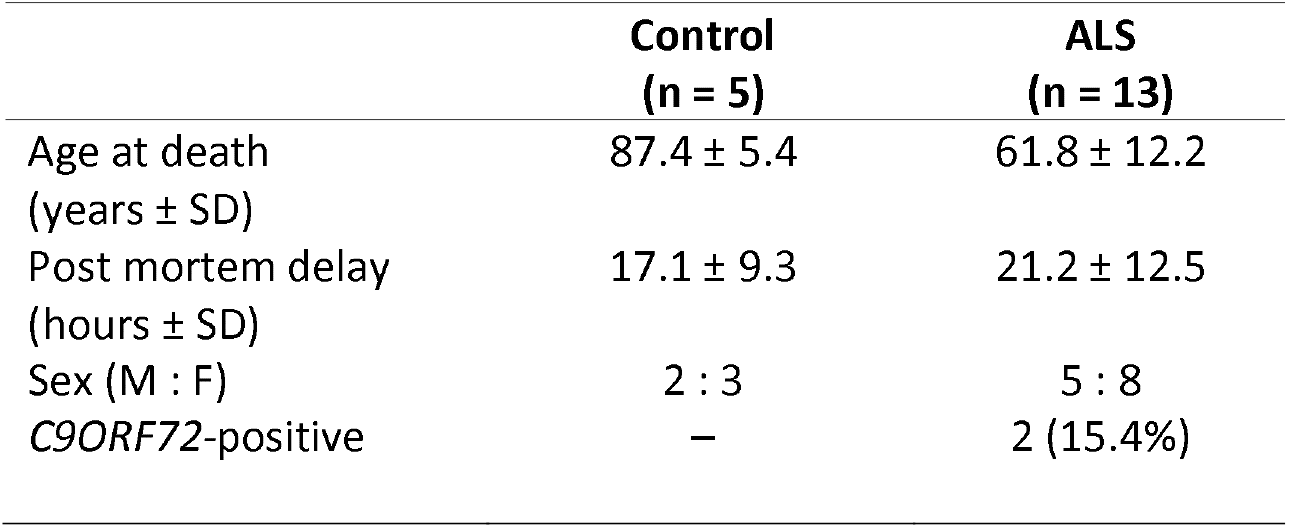
Post mortem tissue donor demographics.

### CSF hemoglobin and total protein measurements

CSF samples were thawed on ice immediately prior to use, and free hemoglobin levels measured using a human hemoglobin ELISA kit, according to manufacturer’s instructions (Cat# E80-136, Bethyl Laboratories). Briefly, 100 μL of capture antibody (1:100 dilution) was added to each well of a 96-well plate and incubated overnight at 4 °C. The plate was then washed and 100 μL of CSF (diluted 1:10 in blocker casein (Thermo Scientific) in Tris-buffered saline) was added to each plate well for 1 h at room temperature. The plate was washed and 100 μL of detection antibody (1:40,000 dilution) added to each well for 1 h at room temperature. After final washes, 3,3’,5,5’-tetramethylbenzidine substrate solution was added to each well for 15 min in the dark at room temperature, and the reaction quenched by adding 100 μL of 1N HCl. The absorbance was measured at 450 nm using a plate reader. All samples were measured in duplicate, and the final free hemoglobin concentration calculated using a standard curve generated using serially diluted pure human hemoglobin calibrator standard. The limit of detection was 6.25 ng/mL. Total protein concentration in 200 μL CSF was measured using the BCA assay (Thermo Scientific; Rockford, IL).

### Fluorescent immunohistochemistry

Serial spinal cord tissue sections from paraffin-embedded blocks were cut with a microtome in the transverse plane at a thickness of 10 μm and mounted onto Superfrost Plus microscope slides (Thermo Fisher). Following desiccation for a minimum of 1 week at room temperature, three sections per block, each separated by 100 μm (ie. every 10^th^ section) in order to sample different motor neurons, were processed for immunohistochemistry. Slides were heated to 60 °C for 1 h on a hot plate before being dewaxed through an alcohol-zylene-water series: 100% xylene, 2x 30 min; 100% ethanol, 2x 15 min; 95%, 85%, 75% ethanol, 2 min each; water 3x 5 min. Heat-mediated antigen retrieval was performed by immersing slides in sodium citrate buffer pH 6 (Abcam) or Tris-ethylenediaminetetraacetic acid (Tris-EDTA) buffer pH 9 (Abcam) for 2 h in a pressure cooker, followed by 1x phosphate-buffered saline (PBS) washes, 3x 5 min, and drawing of wax borders. Sections were permeabilised with PBS-T (0.1% Triton X-100) for 15 min at 4 °C followed by PBS washes 3x 5 min and blocking in 10% normal donkey serum (NDS) for 1 h. Primary antibodies (Claudin-5, Abcam Ab131259, 1:1000; Collagen IV, Abcam Ab6586, 1:500; Hemoglobin, R&D Systems G-134-C, 1:300; P62, BD Biosciences 610833, 1:200; P-glycoprotein, Abcam Ab170904, 1:125; Phospho-TDP-43, Cosmo Bio TIP-PTD-P02, 1:4000; PolyGA repeat, Proteintech 24492- 1-AP, 1:1000; SMI-32, Covance SMI-32R, 1:800; ZO-1, Invitrogen 33-9100, 1:200) and lectin for blood vessels (Biotin-agglutinin lectin Ulex europaeus, Sigma L8262,1:1000) and were applied overnight in 1% NDS. No primary controls received 1% NDS only. Following washing, secondary antibodies (Invitrogen A11058, A21202, or A21206, 1:500) and Cy5-streptavidin for lectin-labelled blood vessels (Jackson Laboratories 016-170-084, 1:500) were applied for 3 h in 1% NDS. Sections were counterstained with Hoechst 33342 nuclear stain for 5 min at 0.5 μg/mL. Where indicated, autofluorescence was quenched with 1x TrueBlack (Biotium) in 70% EtOH for 30 s at room temperature. Sections were coverslipped with #1.5 coverslips (Menzel-Glaser) using ProLong Diamond Antifade Mountant (Invitrogen). Staining was performed for one section per spinal cord level per case in a single staining run (1 × 3 × 18 = 54 sections per run). This was then repeated twice more, such that run variability did not influence case comparisons.

### Imaging

Sections co-labelled for hemoglobin, lectin and SMI-32; or hemoglobin, lectin and barrier integrity markers; were imaged across the entire section with a MetaSystems VSlide slide scanner at 10x magnification (0.45 NA), using MetaCyte acquisition and stitching software. The Colibri 2 (Zeiss) LED light source was used to acquire DAPI (excitation band 375/38 nm), AlexaFluor 488 (484/25 nm), and Cy5 (631/22 nm), while the X-Cite light source was used for AlexaFluor 594 (560/40 nm). Phospho-TDP-43 (pTDP-43) pathology in the spinal cord ventral horn was imaged with a Nikon Eclipse NiE microscope with a Nikon DS-Ri2 camera using NIS elements (Nikon, Version 4.20) at 20x magnification (0.50 NA). All sections (ALS and controls) were imaged with the same settings for each staining combination. Vessel-associated pTDP-43 inclusions and DPR inclusions were imaged on an Olympus FV1000 confocal microscope at 60x (1.35 NA) and 40x (1.00 NA) magnification, respectively.

### Image analysis

The overall image analysis pipeline is shown in Fig S2A. SMI-32-immunopositive motor neurons in the spinal cord ventral horn were counted manually using ImageJ (version 1.51p, National Institutes of Health) using the polygon and multipoint tools (two clicks per cell body) based on staining intensity above local background. The number of SMI-32-positive motor neurons was normalised to the area of tissue analysed (ie. ventral horn area). An observer blinded to the disease status of the tissues performed a second count using the same methodology but only counting one of the three sections for each level and each case. There was a strong positive linear correlation between motor neuron counts performed by each rater (Fig S2B, Pearson r = 0.858, *p* < 0.0001).

Phospho-TDP-43 (pTDP-43) pathology was quantified automatically using the “Count Nuclei” module within Metamorph to measure the number of pTDP-43 inclusions that met standardised size and intensity criteria. The number of pTDP-43 inclusions was normalised to the number of motor neurons in the ventral horn area analysed. An observer blinded to the disease status of the tissues validated the methodology with manual counting of pTDP-43 inclusion number in ImageJ based on staining intensity above local background. This showed a strong positive linear correlation with the automated analysis (Fig S2C, Pearson r = 0.964, *p* < 0.0001).

Hemoglobin leakage and density of lectin-positive vessels were quantified in an automated fashion using a custom journal for MetaMorph software (version 7.8.10, Molecular Devices). Briefly, each image was separated into spinal cord grey and white matter using the draw regions tool. For grey matter, the “Cell Scoring” module was used to measure the integrated intensity of hemoglobin staining which met specified size and intensity criteria (relative to background) and which was also within a defined radius of lectin-positive blood vessels that met separate size and intensity criteria (relative to background). Simultaneously, this module counted the number of vessels that met these criteria. For white matter, where vessels are scant, the requirement for the hemoglobin staining to be near to lectin-positive blood vessels was removed. For both grey and white matter the integrated intensity of hemoglobin staining and the number of lectin-positive vessels were normalised to the area of tissue analysed. Automated analyses of hemoglobin leakage were validated by manual grading by an observer blinded to disease status of the tissues. Hemoglobin leakage was graded on composite images of hemoglobin and lectin using a semi-quantitative 3-point grading scale, and showed a positive linear correlation with the automated analysis (Fig S2D, Pearson r = 0.640, *p* < 0.0001).

BSCB integrity markers were studied within six of the seven cases with the highest overall hemoglobin leakage (as per Fig 5N) and these markers were quantified in an automated fashion using a custom MetaMorph journal. Briefly, adaptive threshold analysis was applied to identify areas of hemoglobin leakage and the average intensity of the barrier marker was measured separately in white and grey matter within leaked and non-leaked areas of the spinal cord.

### Statistical analysis

Statistical analyses were conducted using GraphPad Prism (version 8.0.2). Statistical significance was set at *p* < 0.05. Control and ALS CSF hemoglobin levels, total protein, and hemoglobin normalised to total protein were compared using the Mann-Whitney test. All pooled comparisons between control and ALS spinal cord tissues were performed using Student’s t-test with Welch’s correction. Comparisons between control and ALS cases at each level of the spinal cord were performed using two-way ANOVA with Sidak’s post-test. Comparisons between spinal cord segmental levels within control and ALS were performed using two-way ANOVA with Tukey’s post-test. Comparisons across the spinal cord for ALS alone were performed using one-way ANOVA with Tukey’s post-test. Comparisons between leaked and non-leaked regions in grey and white matter were performed using repeated measures (both factors) two-way ANOVA with Geisser-Greenhouse correction and Sidak’s post-test. Graphs depict individual cases as points (mean of 3 tissue sections per case), overlaid with mean ± standard error of the mean (SEM) for all cases, unless specified otherwise. Using Grubbs’ test (alpha 0.2), one control case was identified as an outlier for motor neuron number and hemoglobin staining and was thus excluded from all graphical and statistical analyses.

## Results

### Hemoglobin levels in the CSF in living ALS patients

In our studies for ALS biomarkers, we banked cerebrospinal fluid (CSF) from individuals living with ALS (sporadic or familial, n= 236) and healthy controls (n= 87) using standard operating procedures established by the Northeast ALS Consortium (NEALS). We routinely measure hemoglobin levels by ELISA in these samples in order to assess potential blood contamination of the CSF during the lumbar puncture. However, we noted that samples with very high hemoglobin were predominantly from ALS patients, leading us to examine whether hemoglobin itself may be a disease biomarker. Indeed, we saw a significant elevation in CSF hemoglobin levels in patients diagnosed with ALS compared to healthy controls (Median: Control, 39.86 ng/mL; ALS 74.25 ng/mL. Upper quartile: Control, 310.4 ng/mL; ALS, 889.8 ng/mL, Fig 1A, *p* = 0.0043). This was not due to increased total protein in the CSF (Median: Control, 741.0 ng/mL; ALS 778.5 ng/mL, Fig 1B, ns), such that CSF hemoglobin normalised to total protein was still increased in ALS patients compared to controls (Median: Control, 0.056; ALS 0.089. Upper quartile: Control, 0.416; ALS, 1.184, Fig 1C, *p* = 0.0067). These data indicate that hemoglobin, which should be restricted to the intravascular compartment, may leak across the blood-brain, blood-spinal cord, or blood-CSF barrier into the CSF compartment during life.

**Figure 1.**
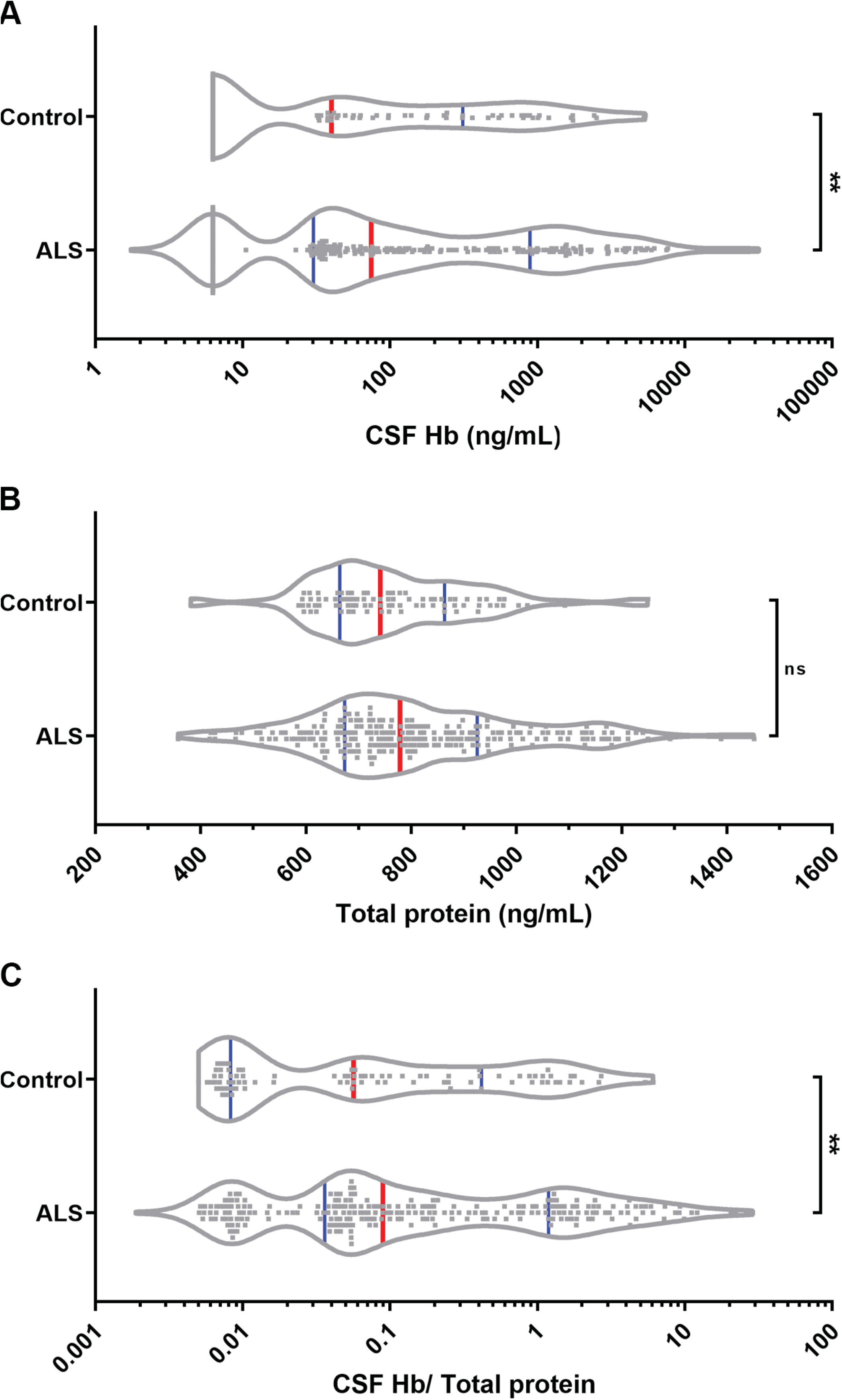
CSF hemoglobin levels in controls and subjects living with ALS. Violin plots (frequency distribution) of CSF from patients with ALS (n= 236) compared to controls (n= 87). Red bars, median; blue bars, quartiles. A) CSF hemoglobin analysis by ELISA. Median: Control, 39.86 ng/mL; ALS 74.25 ng/mL. Upper quartile: Control, 310.4 ng/mL; ALS, 889.8 ng/mL, *p* = 0.0043. B) CSF total protein analysis by BCA, Median: Control, 741.0 ng/mL; ALS 778.5 ng/mL, ns. C) CSF hemoglobin normalised to total protein, Median: Control, 0.056; ALS 0.089. Upper quartile: Control, 0.416; ALS, 1.184, *p* = 0.0067. Mann-Whitney tests.

### Cervical and lumbar enlargements showed reduced transverse diameters in ALS spinal cord

We next utilized post-mortem tissues to examine the relationship between barrier leakage and pathological features along the spinal cord axis in ALS. We required tissue blocks corresponding to the same spinal cord segmental level across a cohort of control and ALS cases. While the majority of our cohort spinal cords were dissected and labelled according to neuronal segmental level of the cord, all controls and several ALS cords were dissected according to bony vertebral segmental level, therefore we first measured and plotted the transverse diameters of all spinal cord segments for each case. For cases cut according to vertebral level, we then employed a conversion scheme to relabel the segments according to neuronal level [30]. After relabelling (Fig 2A,B), the average transverse diameter for control and ALS spinal cords peaked at C5, but ALS cords were narrower at C5 (diameter 9.80 ± 0.43 mm) than controls (diameter 11.71 ± 0.23 mm, Fig 2B, *p* = 0.022). Transverse diameters for the lumbar region of interest L4 were also reduced in ALS cases (diameter 6.82 ± 0.36 mm) compared to controls (diameter 8.51 ± 0.25 mm, Fig 2B, *p* = 0.049).

**Figure 2.**
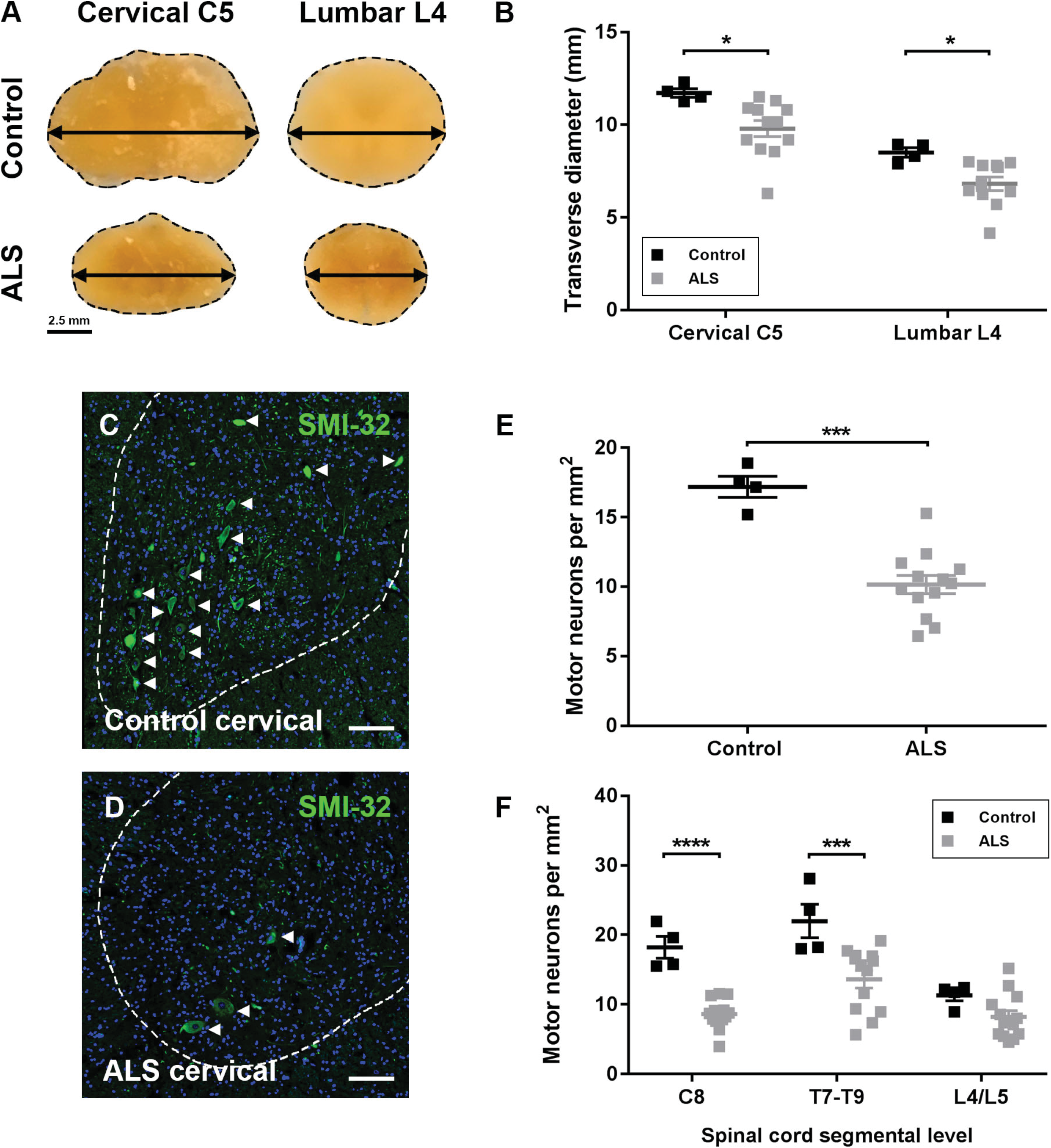
Transverse diameters and motor neuron counts in control and ALS spinal cords. A) Photomicrographs of representative paraffin-embedded control and ALS spinal cord segments. Scale bars 2.5 mm. B) Transverse diameters of cervical segment C5 and lumbar segment L4 for control and ALS cases. Cervical C5: Control, 11.71 ± 0.23; ALS, 9.80 ± 0.43 mm (*p* = 0.022). Lumbar L4: Control, 8.51 ± 0.25; ALS, 6.82 ± 0.36 mm (*p* = 0.049). Two-way ANOVA with Sidak’s post-test. C,D) Immunostaining of spinal cord motor neurons with SMI-32 (green) and nuclei with Hoechst 33342 (blue) in control (C) and ALS (D) cervical spinal cord. Arrow heads depict motor neurons. Scale bars 100 μm. E) Motor neuron numbers per area of ventral horn averaged across cervical C8, thoracic T7-T9, and lumbar L4/L5 for control and ALS spinal cords. Control, 17.18 ± 0.76; ALS, 10.15 ± 0.65 motor neurons per mm^2^ (*p* = 0.0001). Student’s t test with Welch’s correction. F) Motor neuron numbers per area of ventral horn at individual segmental levels C8, T7-T9, and L4/L5. Cervical level C8: Control, 18.22 ± 1.56; ALS, 8.62 ± 0.59 motor neurons per mm^2^ (*p* < 0.0001). Thoracic level T7-T9: Control, 21.99 ± 2.42; ALS, 13.63 ± 1.24 motor neurons per mm^2^ (*p* = 0.0003). Lumbar level L4/L5: Control, 11.33 ± 0.81; ALS, 8.21 ± 0.90 motor neurons per mm^2^ (ns). Two-way ANOVA with Sidak’s post-test.

### Motor neuron loss is most severe in the cervical ALS spinal cord

Having verified that ALS spinal cord enlargements were reduced in size macroscopically, we next quantified changes in the numbers of spinal cord motor neurons along the spinal cord axis. Immunolabelling for the neurofilament heavy chain protein using the SMI-32 antibody identified neuronal somata and processes throughout the grey matter, including motor neurons of the ventral horns (Fig 2C, D). Manual counting specifically in the ventral horn revealed significant motor neuron loss in the ALS spinal cord (10.15 ± 0.65 motor neurons per mm^2^) compared to controls (17.18 ± 0.76 motor neurons per mm^2^, Fig 2E, *p* = 0.0001). Motor neuron loss was observed across all three levels of the cord examined; with statistically significant loss compared to controls at C8 and T7-T9 (Fig 2F). At cervical level C8, ALS spinal cords averaged 8.62 ± 0.59 motor neurons per mm^2^, compared to 18.22 ± 1.56 motor neurons per mm^2^ in controls (*p* < 0.0001). At thoracic level T7-T9, ALS spinal cords averaged 13.63 ± 1.24 motor neurons per mm^2^, compared to 21.99 ± 2.42 motor neurons per mm^2^ in controls (*p* = 0.0003). Motor neurons in the ventral horn were not significantly decreased at L4/L5 in ALS (8.21 ± 0.90 motor neurons per mm^2^) compared to controls (11.33 ± 0.81 motor neurons per mm^2^). Site of symptom onset (upper limb; lower limb; other (respiratory, bulbar, frontotemporal dementia)) did not influence the pattern of motor neuron loss along the spinal cord axis (Fig S3A).

### pTDP-43 pathology is most severe in the cervical and lumbar ALS spinal cord

We next examined the presence and load of the hallmark phosphorylated TDP-43 (pTDP-43) deposits in the spinal cord. pTDP-43 was detected in all 13 ALS spinal cords, in the form of filamentous, round, or neuronal cytoplasmic inclusions (NCI), dystrophic neurites (DN) and glial cytoplasmic inclusions (GCI) (Fig 3A-D). Our automated detection of pTDP-43 was set to count inclusions of 8 μm diameter or greater, which identified the majority of neuronal inclusions but few small glial inclusions (Fig 3E, F). Automated analysis showed that pTDP-43 inclusions were selective for ALS (Fig 3G, *p* = 0.0001), and that pTDP-43 inclusion load in ALS cases was frequently higher in the cervical and lumbar regions, and lower in the mid-thoracic region (Fig 3H, ns), regardless of the site of symptom onset (Fig S3B). However, phospho-TDP-43 inclusion load in individual cases did not predict motor neuron loss (Fig S4A).

**Figure 3.**
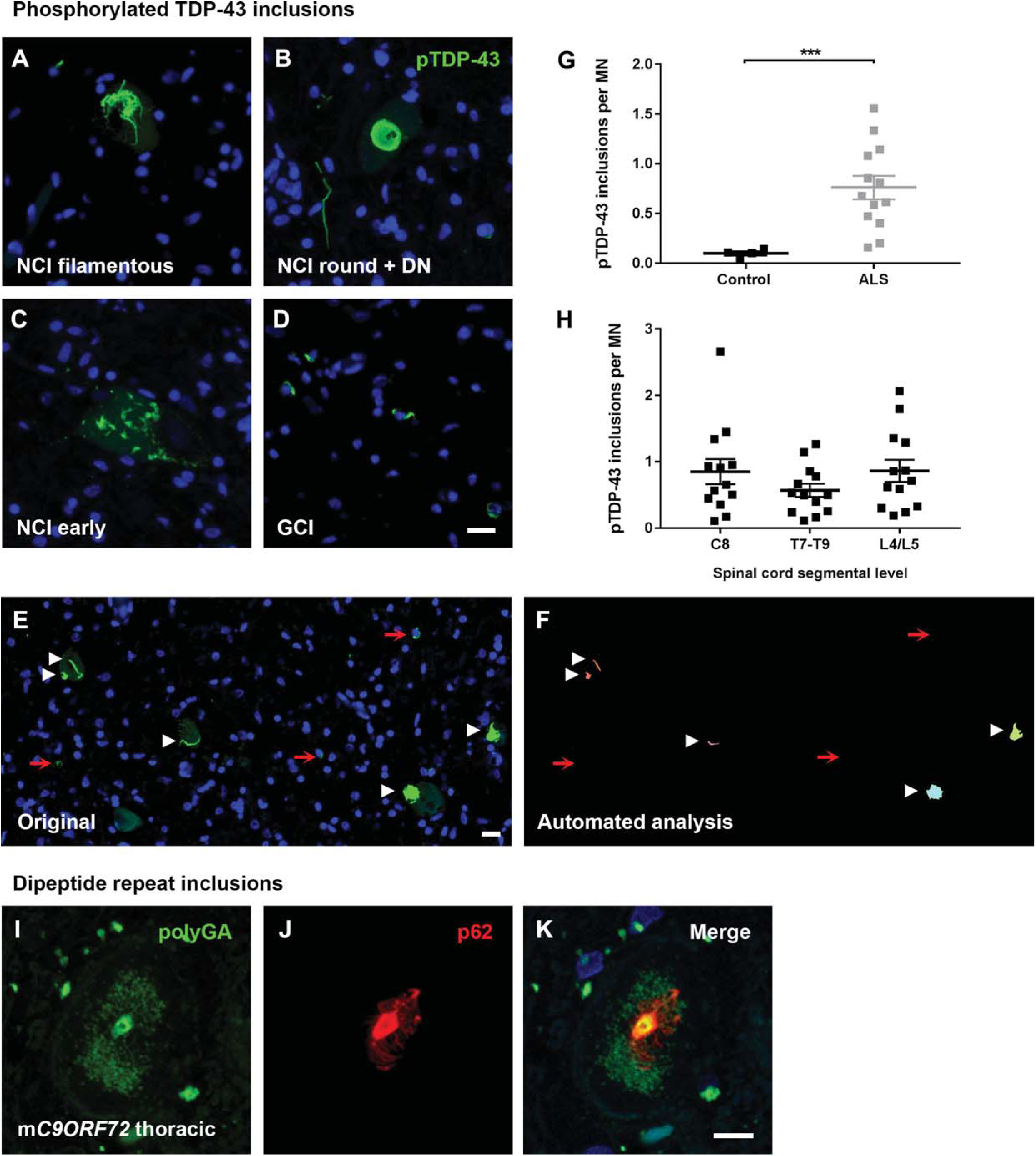
Phosphorylated TDP-43 inclusion pathology across the ALS spinal cord and a rare thoracic polyGA dipeptide repeat inclusion. Immunohistochemical staining for phosphorylated TDP-43 in ALS spinal cords; filamentous (A), round (B), or early (C) neuronal cytoplasmic inclusions (NCI) or dystrophic neurites (DN), and glial cytoplasmic inclusions (GCI) (D). Scale bars 10 μm. E,F) Automated counting of pTDP-43 inclusions of 8 μm diameter or greater detecting neuronal inclusions (white arrow heads) but rarely glial inclusions (red arrows) (E, original image; F, analysis output). G) Phospho-TDP-43 inclusions per motor neuron (Control, 0.10 ± 0.02; ALS, 0.76 ± 0.12, *p* = 0.0001) averaged across cervical C8, thoracic T7-T9, and lumbar L4/L5 for control and ALS spinal cords. Student’s t test with Welch’s correction. H) Phospho-TDP-43 inclusions per motor neuron at individual segmental levels C8, T7-T9, and L4/L5 for ALS spinal cords. Cervical level C8: 0.85 ± 0.19; Thoracic level T7-T9: 0.57 ± 0.10; Lumbar level L4/L5: 0.86 ± 0.17 (ns). One-way ANOVA with Tukey’s post-test. I-K) Confocal maximum projection stack of poly-GA dipeptide repeat inclusions (green) in the thoracic cord of a C9ORF72-positive case, partially colocalised with p62 (red). Scale bars 10 μm.

### PolyGA dipeptide repeat inclusions are rare in C9ORF72-positive ALS spinal cord

In addition to TDP-43 inclusions, other proteins can be deposited in ALS brain and spinal tissue, often depending on the genetic driver of disease. Two ALS cases were known to be positive for the *C9ORF72* repeat expansion mutation, with one case having previously been shown to exhibit abundant polyGP dipeptide inclusions in the hippocampus [31]. We therefore examined the levels of dipeptide repeat protein inclusions along the spinal cord axis. Cervical and thoracic spinal cord sections from both *C9ORF72*-positive cases were stained for the most abundant dipeptide repeat protein (polyGA) [32]. In both cases, polyGA inclusions were extremely rare, and across the four sections studied, only one polyGA inclusion was found that was also positive for the ubiquitin-binding protein p62 (Fig 3I-K). The affected thoracic ventral horn motor neuron showed cytoplasmic colocalisation of polyGA and p62, with neuronal processes embedded in a network of non-colocalised polyGA. Given the dearth of spinal cord dipeptide repeat inclusions, as reported by others [33], we did not map these in finer detail.

### Vessel pathology in ALS spinal cord

Given our interest in the relationship between neurodegeneration, pTDP-43 deposition and integrity of the blood-CNS barriers, we next examined whether pTDP-43 inclusions were found within or associated with the vasculature, and whether the vascular density was normal. While typical parenchymal pTDP-43 pathology was in the form of neuronal or glial cytoplasmic inclusions that were not associated with the vasculature, extremely rare pTDP-43 inclusions were found adjacent to the vessels (Fig 4A-C). These vessel-associated pTDP-43 inclusions were found in three cases with high overall pTDP-43 load; in the cervical cord of two cases, and lumbar cord of one case. We examined the density of vessels in grey and white matter using automated detection and counting of lectin-positive staining (detected as per Fig 4D-G). The density of vessels were higher in ALS than control spinal cords overall, in both grey matter (Fig 4H, p = 0.0041) and white matter (Fig 4I, p = 0.0126), with no differences between segmental levels (Sidak’s post tests, ns).

**Figure 4.**
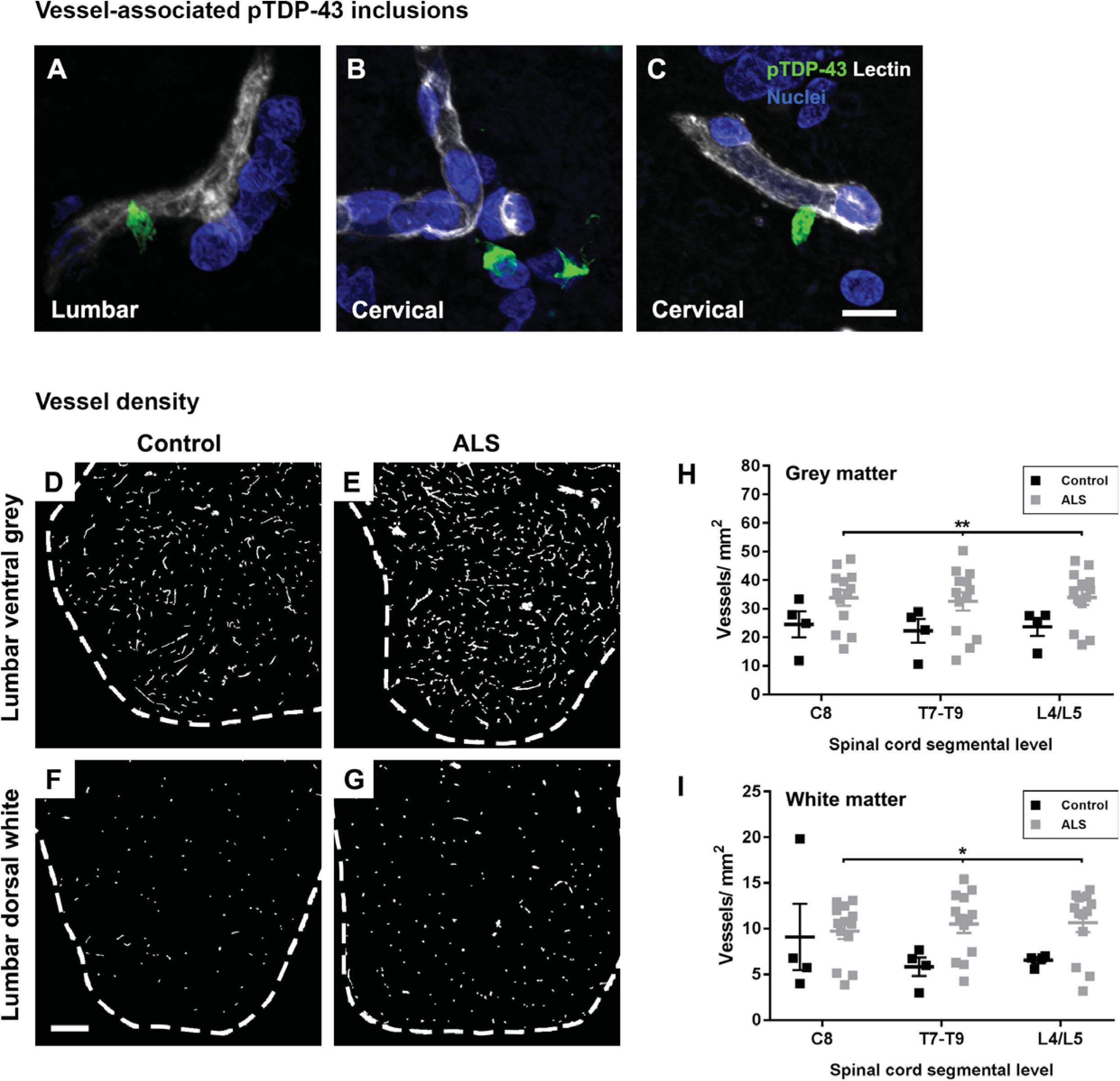
Vessel pathology in ALS spinal cord. A-C) Rare pTDP-43 inclusions (green) adjacent to or associated with lectin-positive blood vessels (white) in ALS lumbar and cervical spinal cord. Scale bars 10 μm. Lectin-positive blood vessel staining in control lumbar spinal cord (D,F) and ALS lumbar spinal cord (E,G). Scale bars 250 μm. H,I) Vessel density at individual segmental levels C8, T7-T9, L4/L5. Grey matter: Cervical level C8: Control, 24.5 ± 4.56; ALS, 33.8 ± 2.76; Thoracic level T7-T9: Control, 22.3 ± 4.12; ALS, 32.6 ± 3.22; Lumbar level L4/L5: Control, 23.7 ± 3.16; ALS, 34.0 ± 2.65 vessels per mm^2^ (overall *p* = 0.0041, post-tests ns). White matter: Cervical level C8: Control, 9.09 ± 3.63; ALS, 9.73 ± 0.87; Thoracic level T7-T9: Control, 5.83 ± 1.02; ALS, 10.50 ± 0.97; Lumbar level L4/L5: Control, 6.54 ± 0.34; ALS, 10.63 ± 1.02 vessels per mm^2^ (overall *p* = 0.0126, post-tests ns). Two-way ANOVA with Sidak’s post-test.

**Figure 5.**
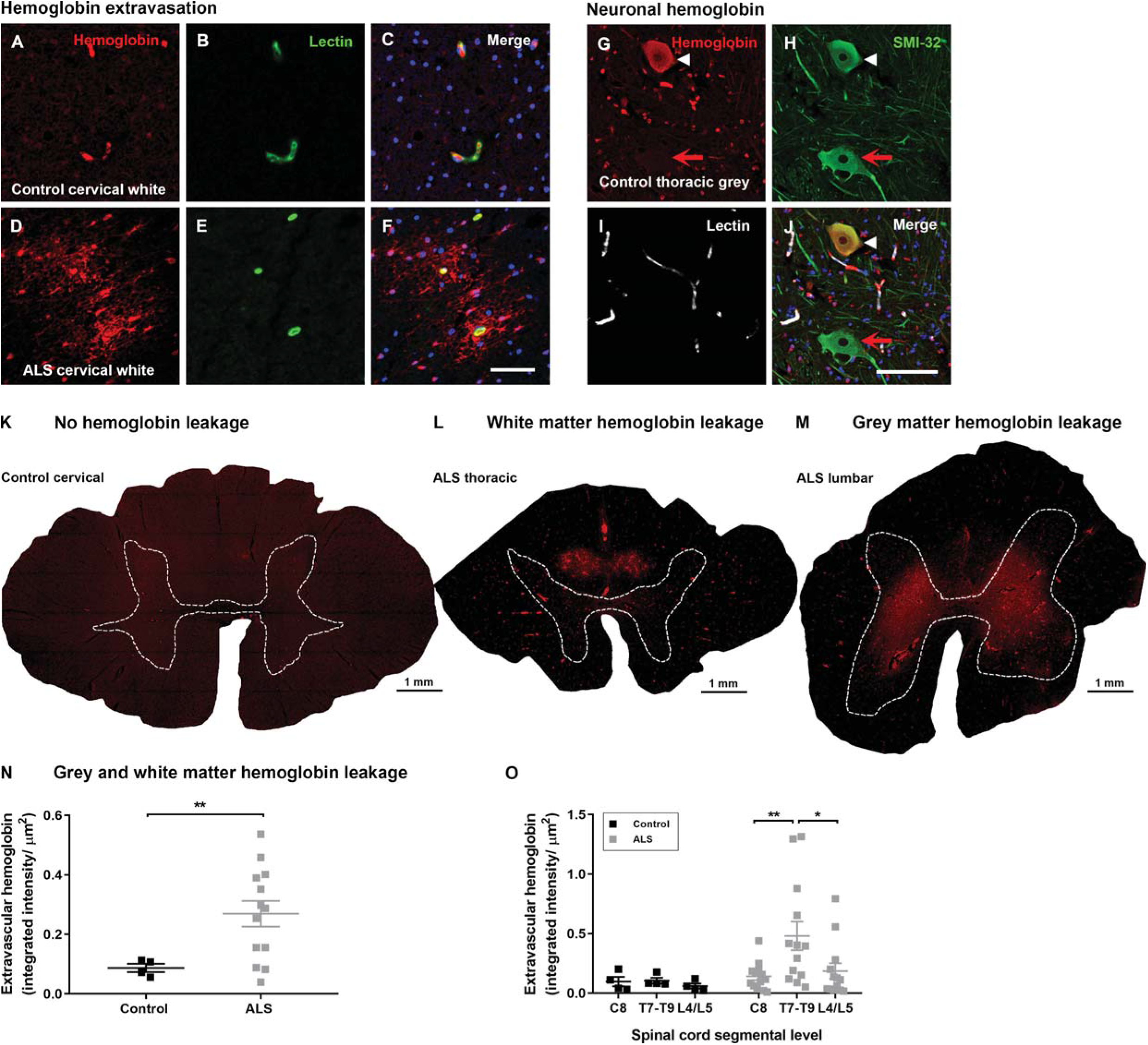
Hemoglobin localisation in control and ALS spinal cord. A-F) Immunohistochemical staining of hemoglobin extravasation in cervical white matter from control (A-C) or ALS (D-F) spinal cord. A, D) Anti-hemoglobin (red); B, E) Lectin-positive vessels (green); C, F) Merged images with Hoechst 33342 (blue). Scale bars 50 μm. G-J) Occasional hemoglobin staining of SMI-32 positive ventral horn motor neurons (compare white arrowhead to red arrow). G) Anti-hemoglobin (red); H) SMI-32-positive motor neurons (green); I) Lectin-positive vessels (white); J) Merged images with Hoechst 33342 (blue), scale bars 50 μm. K-M) Representative images of K) control, L) ALS case with thoracic white matter hemoglobin leakage, M) ALS case with lumbar grey matter hemoglobin leakage (Anti-hemoglobin (red), Lectin-positive vessels (white)). Dashed lines show grey matter boundary. N) Quantification of extravascular hemoglobin in both grey and white matter averaged across cervical C8, thoracic T7-T9, and lumbar L4/L5 levels in control and ALS spinal cord. Controls: 0.087 ± 0.014; ALS: 0.269 ± 0.043 (*p* = 0.0013). Student’s t test with Welch’s correction. O) Quantification of extravascular hemoglobin at individual segmental levels C8, T7-T9 and L4/L5. Controls: Cervical level C8, 0.097 ± 0.039; Thoracic level T7-T9, 0.105 ± 0.024; Lumbar level L4/L5, 0.097 ± 0.0218 integrated intensity (AU) per μm^2^ (ns). ALS: Cervical level C8, 0.481 ± 0.032; Thoracic level T7-T9, 0.185 ± 0.120, Lumbar level L4/L5, 0.185 ± 0.065 integrated intensity (AU) per μm^2^, T7-T9 vs. cervical (*p* = 0.005), T7-T9 vs. lumbar (*p* = 0.017). Two-way ANOVA with Tukey’s post-test.

### Hemoglobin is found in the parenchyma around blood vessels in both grey and white matter ALS spinal cord

We next assessed blood-spinal cord barrier integrity by immunostaining spinal tissue for hemoglobin, a red blood cell protein found previously to leak across the vessel wall into the cervical spinal cord in ALS [20]. While hemoglobin was localised predominantly within blood vessels in control spinal cords (Fig 5A-C), there was extravascular hemoglobin staining in most ALS spinal cords (Fig 5D-F). Extravascular hemoglobin in ALS cases showed a radial distribution around blood vessels, implicating vessel leakage as its source (Fig 5F). In white matter, hemoglobin leakage was seen focally in the dorsal aspects (Fig 5L) and was present in 12/13 ALS cases (92%), while in grey matter the pattern of leakage was diffuse and mostly dorsal of the transverse midline (Fig 5M) and was observed in 10/13 ALS cases (77%). No hemoglobin leakage was evident in controls, although background staining intensity tended to be high (Fig 5K). In some control and ALS cases, neuronal hemoglobin staining was observed in neurofilament H-positive ventral horn motor neurons (Fig 5G-J), as is seen in human cortical neurons (Richter et al., 2009; Codrich et al., 2017), or in cells in the dorsal white matter of the spinal cord which were negative for markers of microglial (Iba1, CD14) or astrocytes (GFAP) (images not shown). Hemoglobin load in individual cases was not correlated with duration of disease (Fig S4B) or post mortem delay (Fig S4C).

### Hemoglobin leakage is differential along the ALS spinal cord axis being most severe at the mid-thoracic level

We next conducted automated image analysis of hemoglobin leakage. The integrated intensity of leaked hemoglobin, averaged across the three studied levels of the spinal cord for each case, was significantly higher in ALS spinal cords than in controls (Fig 5N, *p* = 0.0013). However, because of the finding of higher background staining for hemoglobin in controls, which may pertain to the different fixation of control and ALS tissues, we also analysed the differences in hemoglobin leakage along the cord within, rather than between, the individual cohorts. Comparing the three spinal cord levels within either control or ALS cohorts revealed uniform extravascular hemoglobin at all three levels in controls (Fig 5O, ns), whereas patterning was observed along the spinal cord axis in ALS cases, with leakage more severe at the mid-thoracic level than the cervical (*p* = 0.005) or lumbar levels (*p* = 0.017). This pattern was maintained regardless of the site of symptom onset (Fig S3C).

We next examined hemoglobin leakage in the grey and white matter separately, by calling manually drawn image ‘regions’ into our automated analysis to isolate individual grey and white matter images. Grey matter leakage was bilateral and in the mid-dorsal aspect of the grey matter, being mainly absent from the ventral horns, the spinal canal, or laminae I or II (Fig 6A). Grey matter leakage was detected by automated analysis as per Fig 6B. The integrated intensity of leaked hemoglobin staining in the grey matter, averaged across the three studied levels of the spinal cord for each case, was significantly higher in ALS spinal cords than in controls (Fig 6C, *p* = 0.012). Comparing the three spinal cord levels within either control or ALS cohorts showed a low and consistent level of extravascular hemoglobin at all three levels in controls (Fig 6D, ns), whereas ALS cases exhibited more variable grey matter leakage that was highest at the mid-thoracic level (Fig 6D, ns).

**Figure 6.**
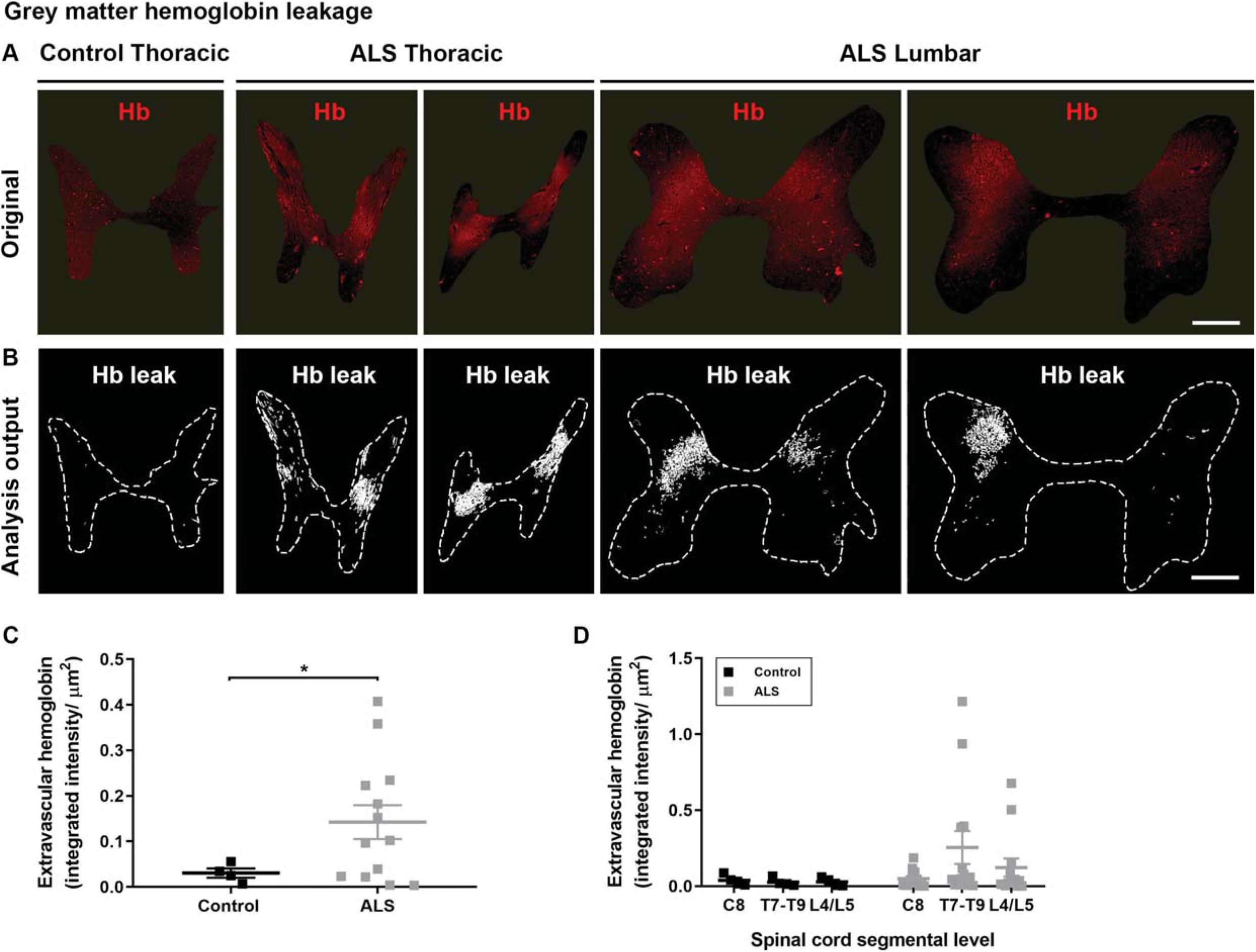
Patterning of grey matter hemoglobin leakage along the spinal cord axis. A) Immunohistochemical staining and B) automated analysis outputs of extravascular hemoglobin in control and ALS spinal cord grey matter. Scale bars 1 mm. C) Quantification of extravascular hemoglobin in grey matter averaged across cervical C8, thoracic T7-T9, and lumbar L4/L5 levels in control and ALS spinal cord. Controls: 0.030 ± 0.010; ALS: 0.142 ± 0.037 (*p* = 0.012). Student’s t test with Welch’s correction. D) Quantification of extravascular hemoglobin in grey matter at individual segmental levels C8, T7-T9 and L4/L5. Controls: Cervical level C8, 0.038 ± 0.018; Thoracic level T7-T9, 0.026 ± 0.014; Lumbar level L4/L5, 0.028 ± 0.014 integrated intensity (AU) per μm^2^ (ns). ALS: Cervical level C8, 0.050 ± 0.015; Thoracic level T7-T9, 0.255 ± 0.109, Lumbar level L4/L5, 0.122 ± 0.060 integrated intensity (AU) per μm^2^ (ns). Two-way ANOVA with Tukey’s post-test.

White matter hemoglobin leakage in ALS spinal cords was predominantly dorsomedial (Fig 7A). White matter leakage was detected by automated analysis as per Fig 7B. The integrated intensity of leaked hemoglobin in the white matter, averaged across the three studied levels of the spinal cord for each case, was significantly higher in ALS spinal cords than in controls (Fig 7C, *p* = 0.0013). Comparing the three spinal cord levels within either control or ALS cohorts showed a low level of extravascular hemoglobin at all three levels in controls (Fig 7D, ns), whereas in ALS cases the overall increase in white matter hemoglobin leakage was shown to be largely driven by the mid-thoracic level, which showed significantly greater leakage than the cervical (*p* = 0.0009) or lumbar levels (*p* < 0.0001).

**Figure 7.**
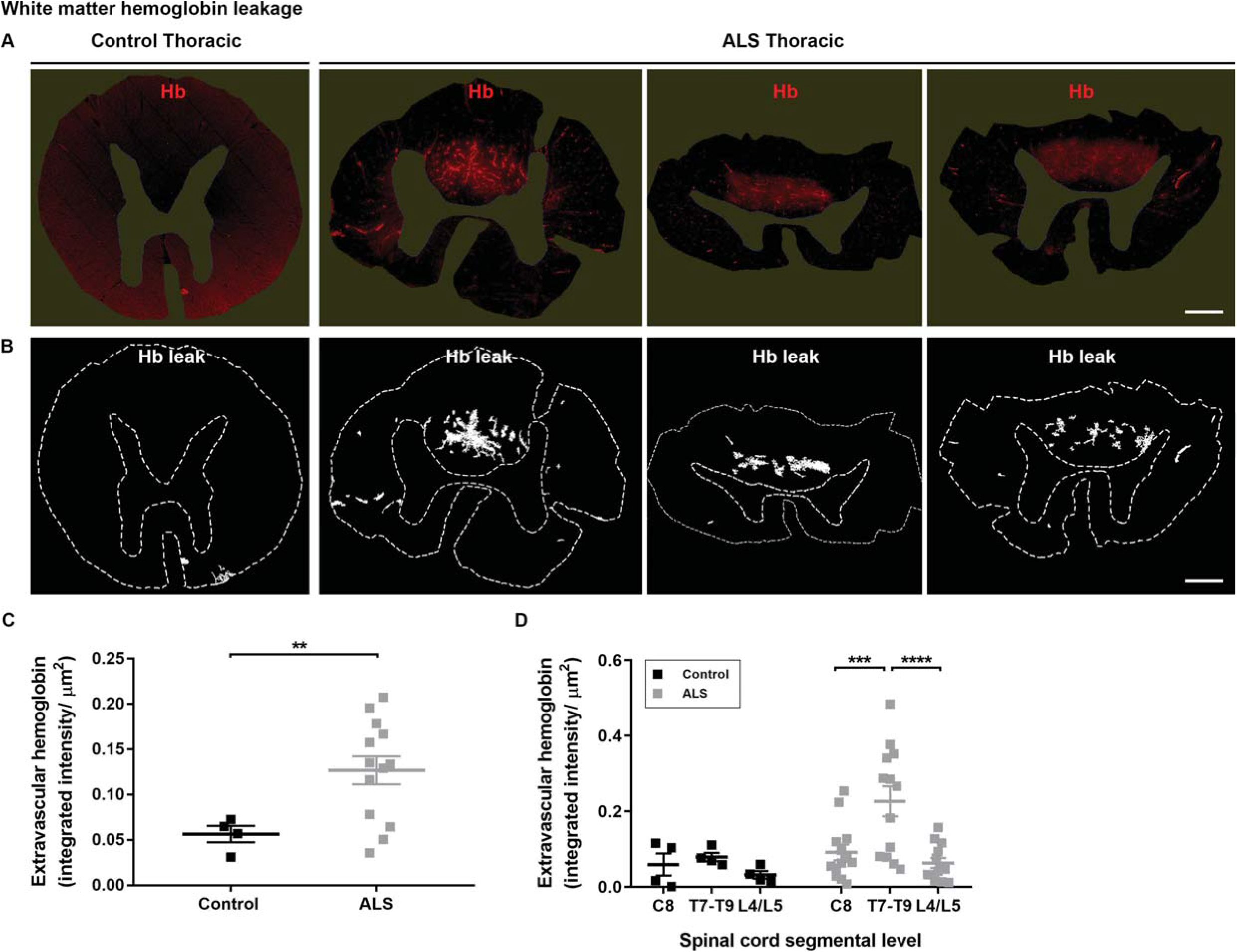
Patterning of white matter hemoglobin leakage along the spinal cord axis. A) Immunohistochemical staining and B) automated analysis outputs of extravascular hemoglobin in control and ALS spinal cord white matter. Scale bars 1 mm. C) Quantification of extravascular hemoglobin in white matter averaged across cervical C8, thoracic T7-T9, and lumbar L4/L5 levels in control and ALS spinal cord. Controls: 0.056 ± 0.009; ALS: 0.127 ± 0.015 (*p* = 0.0013). Student’s t test with Welch’s correction. D) Quantification of extravascular hemoglobin in grey matter at individual segmental levels C8, T7-T9 and L4/L5. Controls: Cervical level C8, 0.059 ± 0.029; Thoracic level T7-T9, 0.079 ± 0.011; Lumbar level L4/L5, 0.032 ± 0.010 integrated intensity (AU) per μm^2^ (ns). ALS: Cervical level C8, 0.091 ± 0.021; Thoracic level T7-T9, 0.226 ± 0.040, Lumbar level L4/L5, 0.063 ± 0.013 integrated intensity (AU) per μm^2^, T7-T9 vs. cervical (*p* = 0.0009), T7-T9 vs. lumbar (*p* < 0.0001). Two-way ANOVA with Tukey’s post-test.

Consistent with hemoglobin leakage occurring in regions distinct from the major motor neuron pathology, hemoglobin load in individual cases did not predict motor neuron number (Fig S4D) or pTDP-43 inclusion load (Fig S4E).

### No loss of selected endothelial and basement membrane markers in areas of hemoglobin leakage

To investigate the mechanisms for pathological passage of hemoglobin across the BSCB in ALS, selected BSCB component proteins were stained in a subset of ALS cases with high hemoglobin leakage. The endothelial tight junction proteins claudin-5 and ZO-1, the efflux transporter P-glycoprotein, and the basement membrane protein collagen IV, were all strongly immunodetected within both leaked and non-leaked patches in the spinal cord, as determined by hemoglobin co-labelling (Fig 8A-H). We used automated detection of patches of hemoglobin leakage (Fig 8N), together with manually drawn grey and white matter ‘regions’, to compare the staining intensity of BSCB components in leaked and non-leaked regions of the spinal cord, in both the grey and the white matter (Fig 8O,P). No changes in staining intensity were observed between leaked and non-leaked areas of the spinal cord, in either grey or white matter, for any of the BSCB components examined (Fig 8I-L, ns).

**Figure 8.**
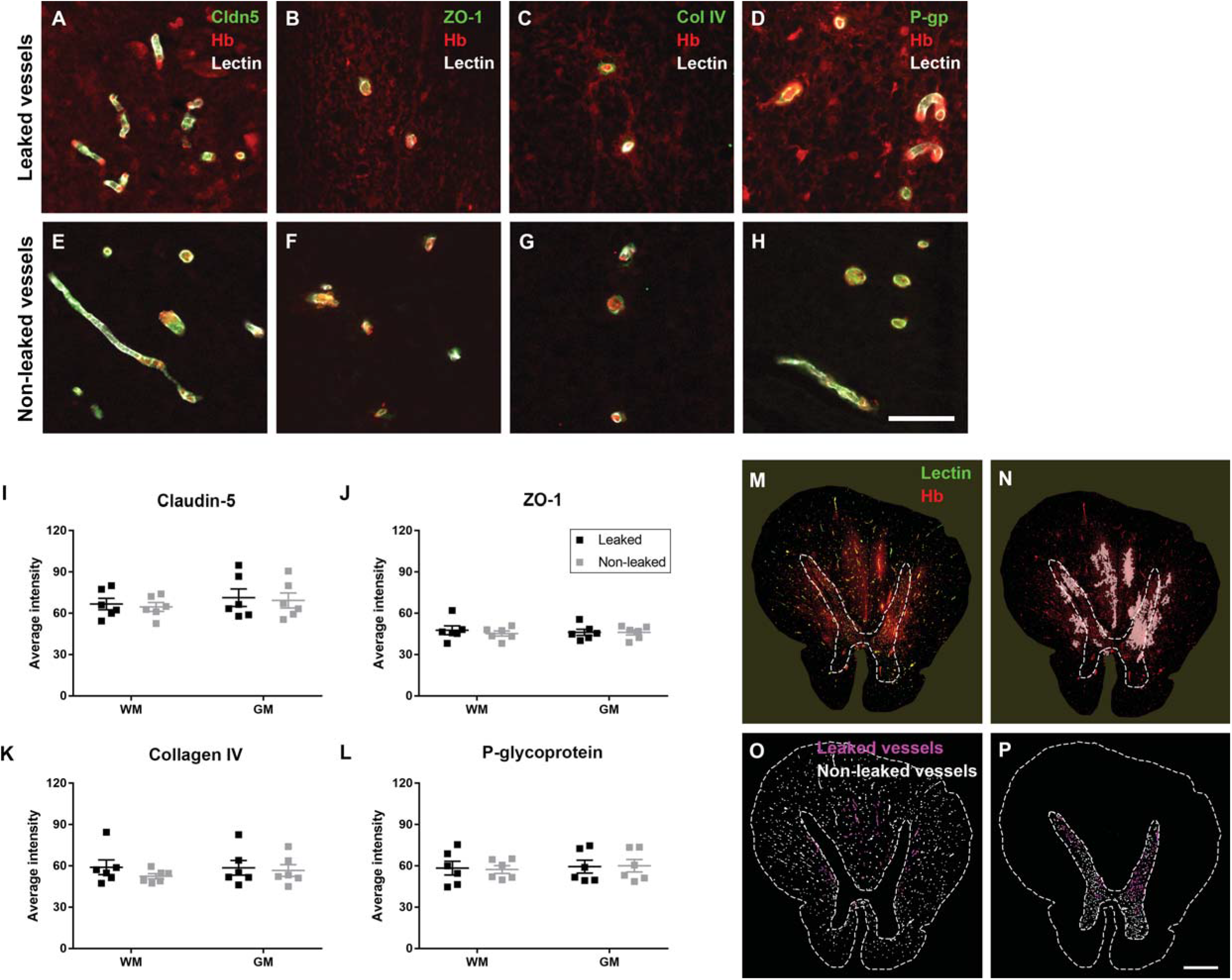
BSCB integrity marker staining between leaked and non-leaked areas of the ALS spinal cord. A-H) Immunohistochemical staining of BSCB integrity markers; tight junctions claudin-5 (A, E) and ZO-1 (B, F), basement membrane marker collagen IV (C, G) and the efflux pump P-glycoprotein (D, H) in spinal cord vessels with (upper) or without (lower) hemoglobin leakage. Scale bar 50 μm. I-L) Quantification of average intensity staining of BSCB integrity markers in a subset of ALS cases with high hemoglobin leakage (n=6) in leaked and non-leaked areas of the white and grey matter of the spinal cord (ns). Two-way repeated-measures ANOVA with Geisser-Greenhouse correction and Sidak’s post-test. M-P) Automated analysis workflow for quantification of BSCB integrity marker staining. Dashed lines show grey matter boundary. Scale bar 1 mm. M) Composite of original images showing anti-hemoglobin (red) and lectin-positive vessels (green). N) Overlay of hemoglobin leakage analysis output (white, partly transparent) over anti-hemoglobin (red). O) Segmentation of vessels inside (magenta) or outside (white) areas of hemoglobin leakage in the white matter. P) Segmentation of vessels inside (magenta) or outside (white) areas of hemoglobin leakage in the grey matter.

## Discussion

Finding high levels of hemoglobin in the CSF of subjects living with ALS, we examined the relationship between blood-spinal cord barrier integrity, endothelial barrier protein expression, motor neuron degeneration and TDP-43 proteinopathy in post-mortem human ALS spinal cord. A key finding of our study was that BSCB compromise marked by hemoglobin deposition in the spinal cord parenchyma around the blood vessels was present along the ALS spinal cord axis. We predicted that BSCB damage would be most severe in the ventral part of the cervical and lumbar cord, given that lower motor neurons in these regions innervate the limbs, which are most commonly affected in ALS [1, 5], and lower motor neuron loss with phosphorylated TDP-43 proteinopathy is greatest in the cervical and lumbar cord [our findings and 5]. Indeed, previous studies found hemoglobin, fibrin, thrombin and IgG extravasation in the cervical spinal cord [20], and fibrin and IgG extravasation in the lumbar spinal cord [21]. The thoracic cord, where the BSCB is ‘tightest’ [34], had not previously been examined. However, we demonstrate that hemoglobin extravasation is also found at the mid-thoracic level, and that this thoracic BSCB leakage is the most severe. The thoracic spinal cord motor neurons innervate the intercostals and some accessory respiratory muscles, and their degeneration is presumed to be a late-stage feature of disease [35]. Intriguingly, we also find that the most striking hemoglobin extravasation does not occur ventrally, adjacent to the spinal motor neurons, but in the dorsal aspects of the grey and particularly the white matter. Our findings therefore call into question the paradigm that motor neuron neurodegeneration induces local barrier leakage.

Lower spinal motor neuron somata are predominately located in lamina IX of the spinal cord in the ventral grey matter, with their axons projecting to target muscle fibres via the ventral roots. Yet leaked hemoglobin was detected across both grey and white matter; most severely in the dorsomedial white matter and rarely in the ventral grey matter. Similarly, microhemorrhages in the cervical and lumbar spinal cord of SOD1 mice were found to be quite uniformly distributed across the grey and white matter, without preference for the ventral grey matter [36]. Therefore, although motor neuron damage has been proposed to induce neuroinflammation and BSCB permeability to infiltrating immune cells [21], our data argue against BSCB changes in ALS being initiated by nearby neuronal damage.

Hemoglobin leakage was only seen in ALS spinal cord tissue in our study, but it should be noted that our control and ALS spinal cords were preserved using different fixatives-Dodge^TM^ embalming fluid for controls and 15% formalin for ALS cases. While all stains tested, including hemoglobin, stained both tissue types, it is conceivable that differential fixation could confound comparisons between these two groups. Certainly, increased hemoglobin leakage in ALS compared to controls has been reported previously for humans and transgenic ALS animal models [11, 20, 37, 38], and is supported by our finding of elevated CSF hemoglobin in living ALS patients. The strength of our study lies in also characterising and quantifying the patterning of BSCB leakage and BSCB component expression across grey and white matter and along the spinal cord axis within individual ALS spinal cords.

Using such within-case analysis we found no change in junctional components that mediate BSCB tightness, ZO-1, claudin-5, collagen IV or P-glycoprotein, between leaked and non-leaked regions of individual ALS cords. Previous studies found changes in these BSCB components between transgenic ALS models or human patients and their respective controls, and it was presumed that BSCB component changes underpinned increased permeability. For instance, in post-mortem ALS patient spinal cord, endothelial cells showed reduced expression of ZO-1 [21, 39], claudin-5 [21], and occludin [39], and detachment of astrocyte end feet [40]. However, there are some disparities in the literature, with other studies showing that tight junctions in the spinal cord in humans [41] and a TDP-43 conditional knockout mouse [42] were morphologically normal. Similarly, basement membrane collagen IV has been reported to both accumulate [21] and decrease [40]. Between-case variability and the difficulty of normalisation within a dramatically altered system make these hypotheses challenging to test, hence our vessel-by-vessel within-case analyses of the relationship between leakage and BSCB components. We show that several key molecules that mediate BSCB integrity are not differentially expressed at the end of life by vessels that are leaky, and nor does leakage involve proteinopathy of the vascular cells. Our ongoing studies seek to test whether vessels with impaired barrier function are invested with fewer pericytes, the mural cells that suppress endothelial transcytosis [43], as shown by others [20, 44].

We found significantly higher levels of hemoglobin in CSF from people living with ALS. This finding alone cannot distinguish between alterations in CSF production, or disruption at the blood-CSF, blood-brain or blood-spinal cord barriers. However, others also report elevated hemoglobin subunits alpha and beta in ALS CSF compared to controls, in the context of both increases and decreases in other CSF proteins [22]. Taken together with our finding of perivascular hemoglobin deposition in the spinal cord parenchyma, it is reasonable to postulate that elevated CSF hemoglobin in ALS derives at least in part from increased permeability of the BSCB, such that interstitial fluid drainage would enrich the CSF with hemoglobin.

We propose that BSCB breakdown occurs independent of motor neuron pathology in ALS. Several rodent models of ALS show BSCB breakdown prior to motor neuron loss [37, 40], or occurring transiently then resolving [45]. Barrier leakage in human Alzheimer’s disease is also independent of either neurodegeneration or the hallmark proteinopathy, although increased BBB permeability predicts early cognitive dysfunction [46]. Whether leaked hemoglobin in the CSF and tissue can induce symptoms in ALS, and if so at what stage in disease, warrants further study. However, the inverse regional relationship seen here between motor neuron pathology and BSCB leakage in ALS support the view that cell-autonomous dysfunction linked to TDP-43 proteinopathy, and <u>not</u> BSCB dysfunction, is the major driver of motor neuron degeneration.

## Conclusions

Together our findings suggest that BSCB breakdown is differential along and across the spinal cord axis, in a pattern that is independent of that of lower motor neuron death or TDP-43 proteinopathy. BSCB leakage is therefore unlikely to be caused by lower motor neuron degeneration, and similarly lower motor neuron degeneration is likely driven predominantly by factors other than BSCB disruption.

## Declarations

Ethics approval and consent to participate Consent for human brain tissue collection and study was obtained from donors and their families prior to death, and ethical approval for this study was obtained from The University of Auckland Human Participants Ethics Committee (#011654). Consent for publication Not applicable.

## Availability of data and materials

The datasets used and/or analysed during the current study are available from the corresponding author on reasonable request.

## Authors’ contributions

CT conducted neuropathological diagnostics; HJW, RLMF, MAC prepared human tissue for study; SW, BVD, YZ, JA conducted experiments; SW, MEVS, NLG, HCM, JA, RB, ELS performed data analysis or designed analysis methods; SW, ELS wrote the manuscript; MAC, MD, ELS conceived of and designed the study. All authors read and approved the final manuscript.

## Acknowledgements

This publication is dedicated to the patients and families who contribute to our research. We thank Marika Eszes at the Centre for Brain Research, University of Auckland, New Zealand; the imaging team at the Biomedical Imaging Research Unit, University of Auckland, New Zealand; the Neurological Foundation of New Zealand for their ongoing financial support of the Human Brain Bank; and the Northeast ALS Consortium (NEALS) Biofluid Repository for providing the CSF samples for this study.

**Authors’ information (optional)**

