## Supplementary material for "Blood-spinal cord barrier leakage is independent of motor neuron pathology in ALS": Fig S

**Supplementary Figures and Figure Legends**

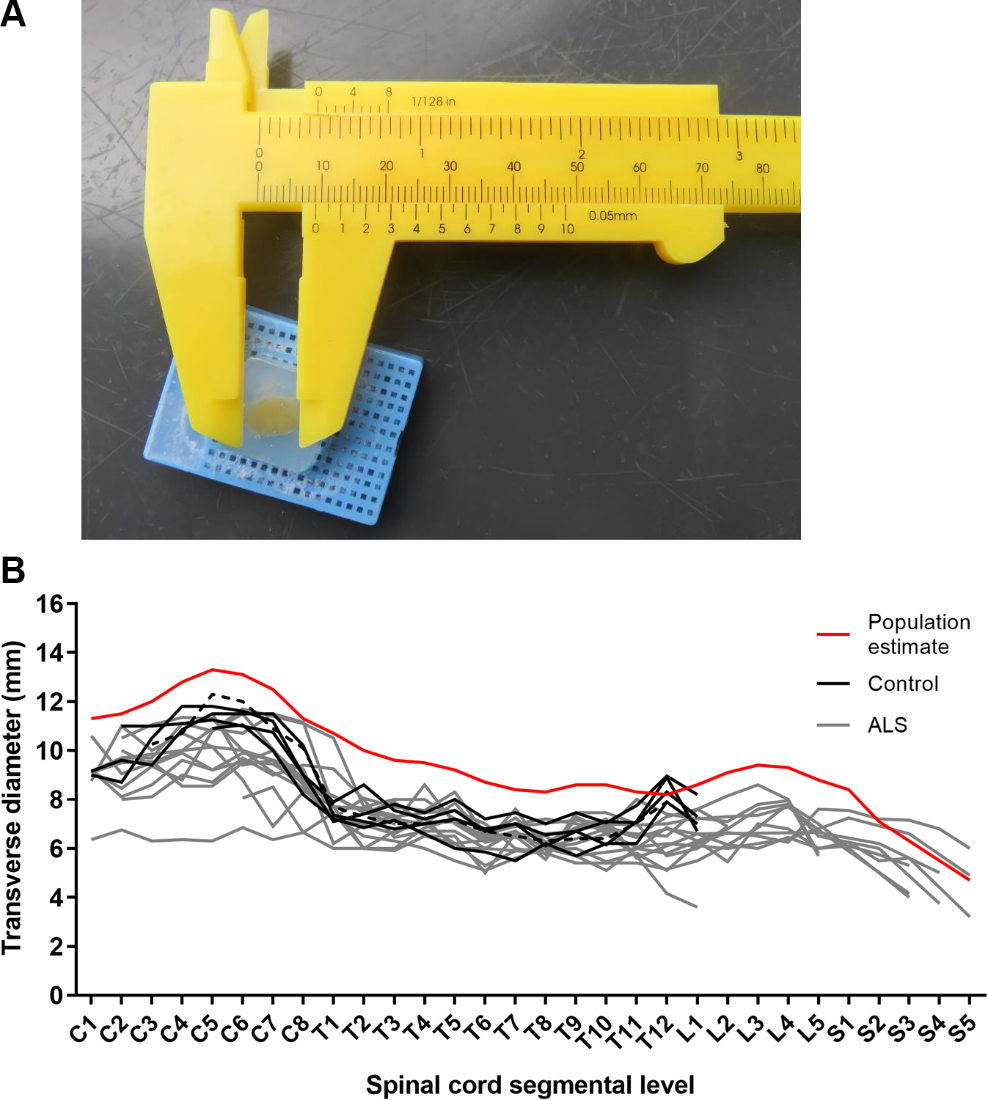

**Figure S1 Spinal cord segment measurement**

A) Image of Vernier callipers being used to measure spinal cord segment transverse diameters (to nearest 0.05 mm). B) Transverse diameters of control and ALS spinal cords prior to conversion, which demonstrates the need for vertebral-to-neuronal segment labelling conversion. Population estimate from [1]. Dashed line indicates the control case that was an outlier for motor neuron counts and hemoglobin staining.

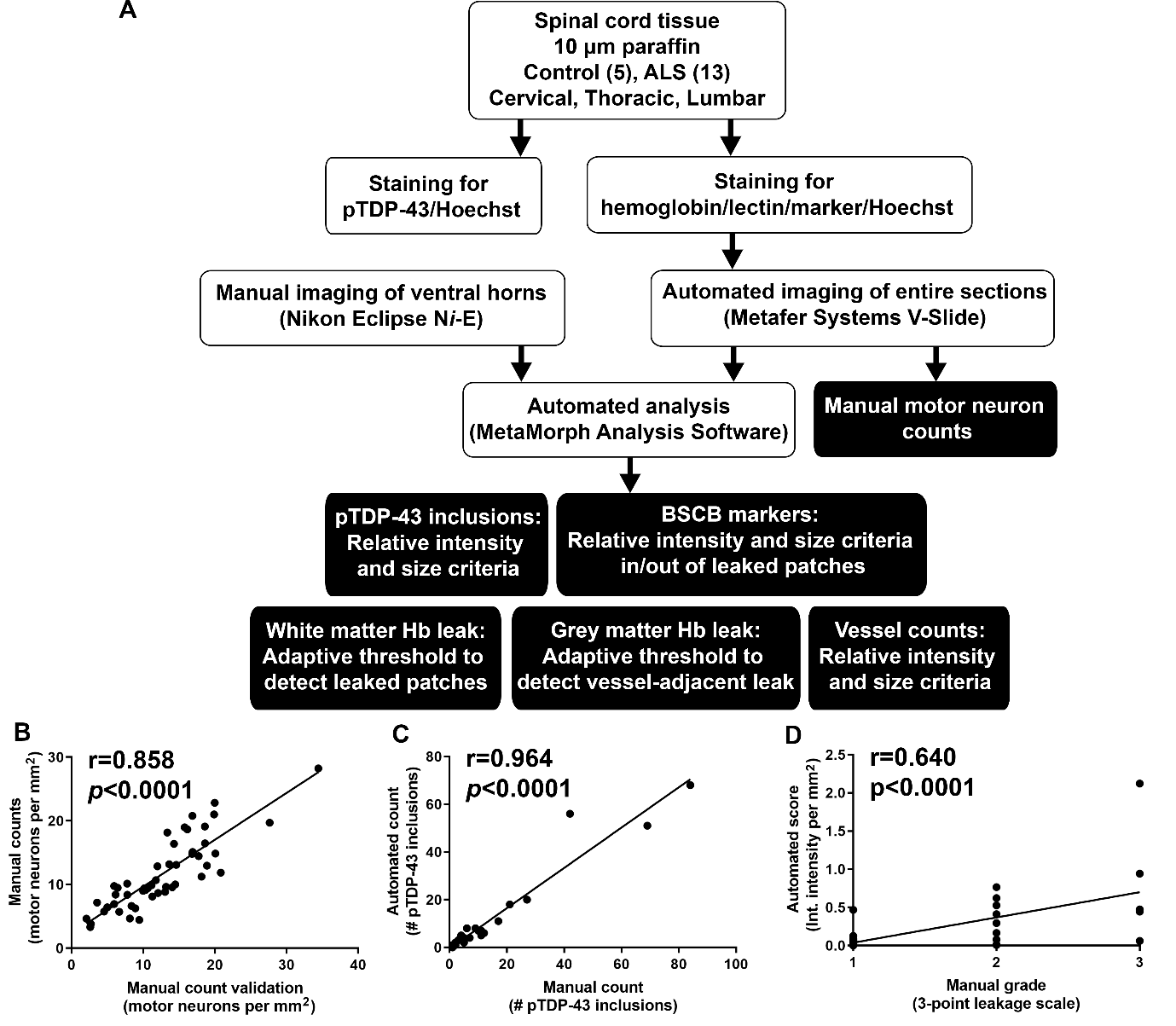

**Figure S2 Tissue processing and analysis**

A) Workflow of tissue staining and quantitative image analysis methodology where 10 µm sections of paraffin-embedded human spinal cord were stained for either pTDP-43 or for hemoglobin and lectin with a marker of interest (SMI-32, P-gp, claudin-5, collagen IV, or ZO-1). Subsequent imaging with the Nikon or VSlide, and manual counting analysis for SMI-32-positive motor neurons, or automated analysis for pTDP-43 pathology, hemoglobin leakage and vessel density. B) Validation of manual motor neuron counting against manual counting of motor neurons performed by a blinded observer, Pearson r=0.858 (p <0.0001). C) Validation of automated pTDP-43 inclusion analysis against manual counting of pTDP-43 inclusions, Pearson r=0.964 (p < 0.0001). D) Validation of automated hemoglobin analysis against manual scoring by a blinded scorer on a semi-quantitative 3-point leakage scoring scale, Pearson r=0.640 (p < 0.0001).

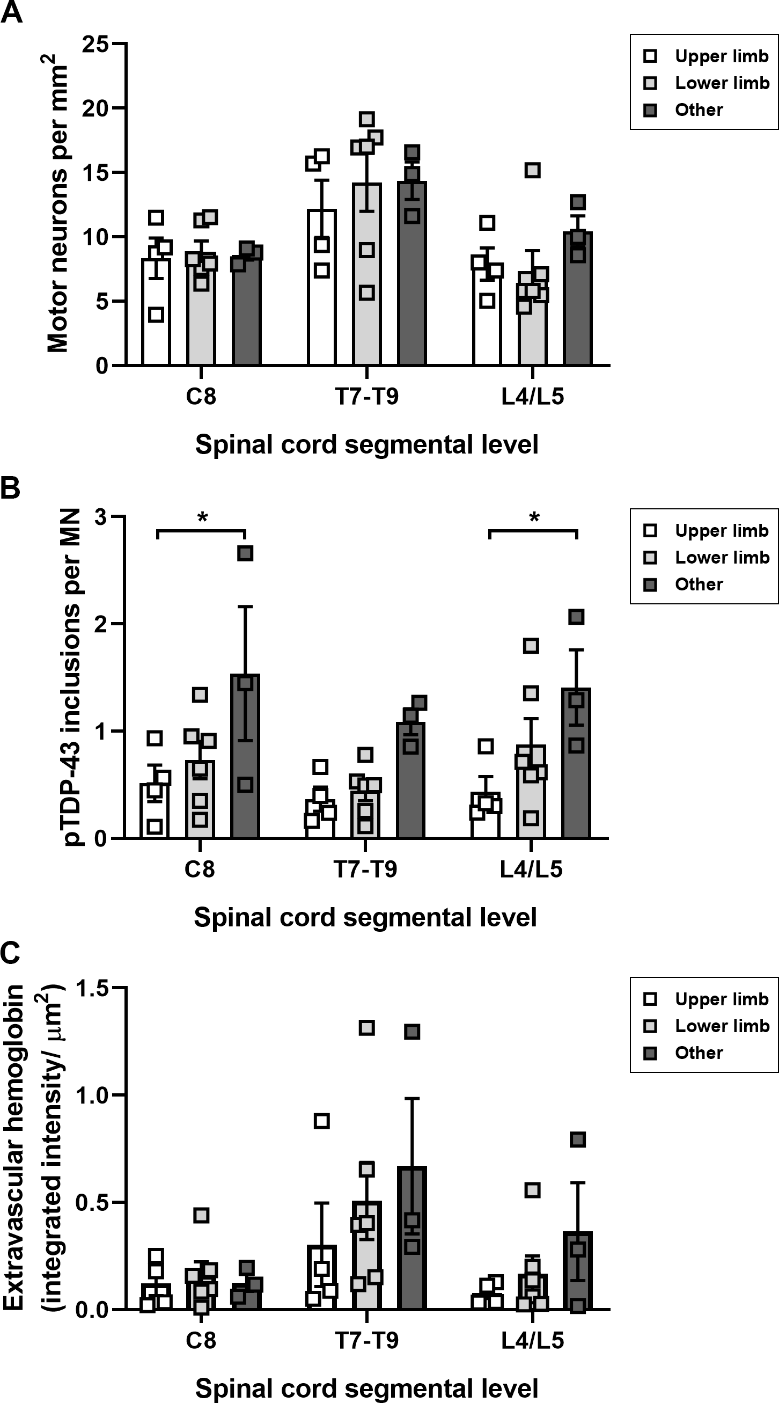

**Figure S3 Pathological measures with respect to site of symptom onset**

A) Motor neuron numbers per area of ventral horn at individual segmental levels C8, T7-T9, and L4/L5 with respect to site of ALS symptom onset; upper limb, lower limb, or ‘other’ (respiratory, bulbar, frontotemporal dementia). B) Phospho-TDP-43 inclusions per motor neuron at individual segmental levels C8, T7-T9, and L4/L5 with respect to site of ALS symptom onset. C8 and L4/L5: Upper limb vs. other (*p* < 0.05). C) Quantification of extravascular hemoglobin (both grey and white matter) at individual segmental levels C8, T7-T9 and L4/L5 with respect to site of ALS symptom onset. Two-way ANOVA with Tukey’s post-test.

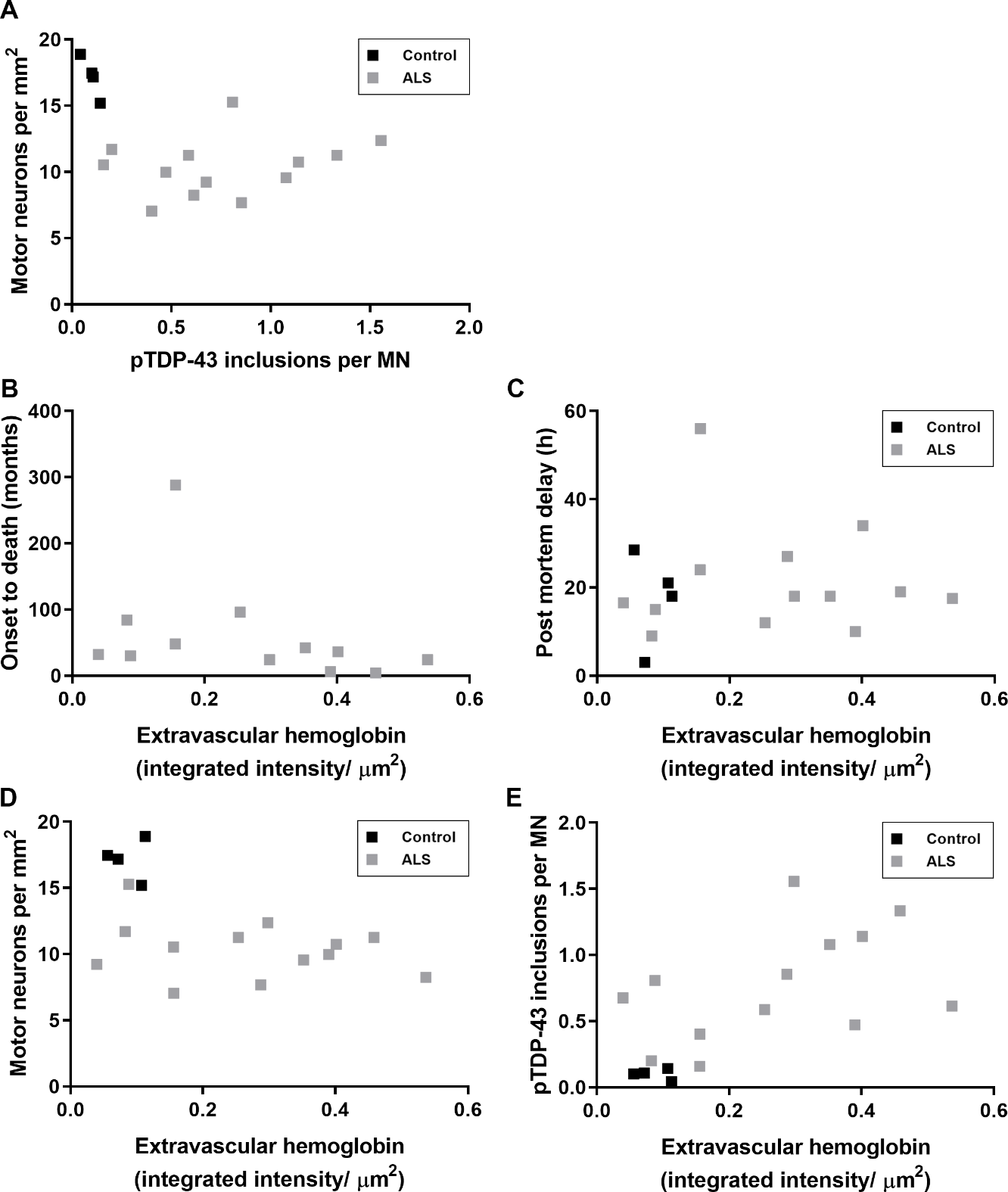

**Figure S4 Extravascular hemoglobin did not correlate with disease, spinal cord tissue collection delay, or motor neuron pathology**

A) Phospho-TDP-43 inclusions per motor neuron did not correlate with motor neuron numbers per area of ventral horn. B-E) Extravascular hemoglobin in both grey and white matter averaged across cervical C8, thoracic T7-T9, and lumbar L4/L5 levels in ALS spinal cord did not correlate with B) disease duration, C) post mortem delay, D) motor neuron numbers per area of ventral horn, or E) phospho-TDP-43 inclusions per motor neuron. Pearson correlation conducted on ALS cases only, all ns.
